# Tumor-associated Macrophages protect Glioblastoma cells from ferroptosis by inducing the release of Ferritin-bound iron via exosomes

**DOI:** 10.1101/2025.10.21.683804

**Authors:** Aurosman Pappus Sahu, Kondaiah Palsa, Ganesh Shenoy, Becky Slagle-Webb, James R Connor

## Abstract

Tumor-associated macrophages (TAMs) are the most abundant non-tumor cell type in glioblastoma (GBM) and act as the pivotal cell type in regulating iron metabolism in the GBM tumor microenvironment. High TAM infiltration into the TME is also associated with increased resistance to ferroptosis, an iron-dependent cell death. However, the exact mechanism by which TAMs make the cancer cells resistant to ferroptosis remains relatively unexplored. Here, we have investigated how TAMs modify iron metabolism in GBM cells to make them more resistant to ferroptotic stress. We utilized GL261 cells, a GBM cell line derived from C57BL6 mice, and syngeneic primary murine bone marrow-derived macrophages (BMDM) to study GBM-TAM interactions in vitro. We found that male macrophages exhibited higher iron uptake, greater iron storage, and a larger labile iron pool compared to female macrophages, indicating intrinsic sex biases in macrophage iron metabolism. Subsequently, we used co-culture experiments to study how macrophages regulate the iron and ferroptotic status of GL261 cells. We discovered that GL261 cells cocultured with BMDMs showed higher resistance to RSL3-induced ferroptotic stress. Mechanistically, BMDMs caused a decrease in total cellular iron in GL261 cells by inducing increased H-ferritin-bound iron release via CD63-positive exosomes; thus, limiting the amount of iron that is available for lipid peroxidation during ferroptosis. This process was moderately sex biased in favor of male macrophages. Finally, we show that this mechanism of BMDM-induced resistance to ferroptosis is independent of Hepcidin regulation and can act as a possible pathway by which GBM cells escape ferroptotic stress during proinflammatory conditions.

## Introduction

Glioblastoma (GBM) is one of the most difficult-to-treat cancers, with a 5-year survival rate of around 6 percent [1]. It is a solid tumor of astrocytic origin, which is characterized by a unique tumor microenvironment (TME) consisting of tumor cells, brain microglia, and infiltrating immune cells [2]. Among the infiltrating cells, monocytes are known to migrate into the tumor, where they subsequently differentiate into tumor-associated macrophages. TAMs comprise up to 50 percent of the total tumor volume and are one of the most abundant cell populations in the TME. The role of TAMs in tumorigenesis has been extensively documented, especially in terms of cytokine secretion, immune cell recruitment, and altering the metabolic landscape of the TME [3]. Especially in iron metabolism, TAMs play a significant role in maintaining iron homeostasis in the TME and can play a pivotal role in controlling ferroptotic stress in neoplastic cells [4].

The role of iron in tumor progression has been understudied, especially in the context of GBM [5, 6]. Iron plays a vital role in human physiology, given that it is actively involved in oxygen transport. Iron is essential for several cellular processes, such as mitochondrial ATP production, DNA synthesis, and cell proliferation [7-9]. Cancer cells have a higher iron requirement than normal cells; hence, they upregulate the expression of transferrin receptor (TFRC), a protein involved in iron uptake, and ferritin, a protein involved in cellular iron storage. In GBM, particularly, both transferrin receptor and ferritin are necessary for tumorigenesis [10]. However, due to the high iron levels found in cancer cells, they are susceptible to ferroptosis, an iron-mediated form of cell death. Ferroptosis is characterized by increased free iron and lipid peroxidation [11]. Lipid peroxidation occurs as a result of the Fenton reaction, where ferrous iron (Fe^2+^) reacts with H_2_O_2_, producing hydroxyl radicals. These hydroxyl radicals further react with phospholipid hydroperoxides to form phospholipid hydroperoxide radicals, which can cause membrane damage [12]. Ferroptosis depends primarily on the ferrous (Fe^2+^) iron in the cytoplasm (labile iron pool), and protein-bound ferric iron (Fe^3+^, stored within ferritin) is not directly involved in ferroptosis. As a result, ferritin can act as a ferroptosis suppressor by quenching available Fe^2+^ in the labile iron pool [13].

In several tumor types, it has been reported that TAMs can affect the ferroptotic state of the tumor cells. This is because TAMs regulate how much iron is available to the cancer cells by direct iron delivery, as well as tumor iron retention and release by different signaling pathways. The only known mechanism by which TAMs protect tumor cells from ferroptosis is by facilitating FPN-mediated iron export from cancer cells, resulting in ferroptosis suppression [14]. However, in GBM, due to the proinflammatory nature of the TME, FPN-mediated iron export is inhibited due to the elevated expression of Hepcidin. Hence, there may be alternate ferroptosis suppression pathways active during inflammation that are independent of the FPN/Hepcidin regulation.

GBM is a sexually dimorphic disease, with sexually dimorphic features in incidence, phenotype, response to therapy, and overall outcome [15, 16]. GBM is more frequent and aggressive in male patients. In terms of tumor incidence, males have a 1.6-fold higher chance of developing GBM than females. Males also develop more primary (de novo) tumors, which are characterized by fast-growing tumors without any less malignant precursor lesions. On the other hand, females develop more secondary tumors, which are slow-growing and originate from lower-grade astrocytomas [17]. Females also have longer survival and show better responses to standard therapy [18]. Sex difference in iron metabolism is thought to be a major contributor to the sexually dimorphic nature of GBM [19-22]. Most studies addressing ferroptosis and iron metabolism in glioblastoma focus on a single specific cell line model, underestimating neoplastic cell-TAM interaction and the sexually dimorphic nature of the tumor microenvironment. Here, we use murine primary macrophages and syngeneic GBM cells of both sexes to study how TAMs interact with GBM cells to regulate ferroptosis in the tumor in an in vitro system. In this study, we explore a novel mechanism involving the release of exosomes containing iron-rich FTH1, by which TAMs protect GBM cells from ferroptotic stress. This mechanism, being independent of FPN-mediated iron release, could be essential for protecting GBM cells from ferroptosis during tumor-induced inflammation.

## Methods

### Isolation of primary bone marrow-derived murine macrophages

The isolation of primary macrophages from the bone marrow of C57BL/6 mice was preformed using previously reported protocols. Briefly, femurs and tibias were collected from C57BL/6 mice and flushed with DMEM with GlutaMAX (ThermoFisher Scientific, Catalog #: 61870036) to collect bone marrow. The bone marrow was washed and run through a 70 μm cell strainer and cultured in non-tissue culture-treated Petri dishes in DMEM with GlutaMAX with the addition of 20% v/v conditioned media from L929 cells and 30% v/v FBS to serve as a source of M-CSF for differentiation of bone marrow progenitor cells into macrophages. Additional media was added to the Petri dish on day 3 and day 6. On day 7, the differentiation was confirmed by looking at the cell morphology, adherence to the Petri dish, and expression of macrophage-specific markers (F4/80 and CD11b).

### Cell culture

GL261 cells were cultured in DMEM with GlutaMAX (ThermoFisher Scientific, Catalog #: 61870036) with 10% fetal bovine serum (GeminiBio, Catalog#: 100-106), and 1% Penicillin-Streptomycin (ThermoFisher Scientific, Catalog #: 15140-122). Cells were maintained in a humidified tissue culture incubator with 5% CO2 at 37 °C. For experiments, cells up to passage number 8 post-thawing were used to maintain reproducibility. For the co-culture experiments, the GBM cells were cultured on a 12-well plate, and BMDMs were cultured on transwell inserts of pore size 0.4 µm to facilitate the free exchange of secreted proteins and exosomes.

### RSL3 treatments

RSL3 (Catalog No.S8155, Selleckchem) was used to induce ferroptosis in GBM cells. Briefly, 1 or 3 μM (depending on the experiment) of RSL3-containing media was added to the GBM cells for 24 hours to induce ferroptosis. The expression of GPX4 was confirmed by Western blot to verify RSL3 activity.

### Lipid Peroxidation analysis

Cells were plated in 12 well tissue culture plates and then treated as indicated. The cells were incubated with 5 μmol/L of C11-BODIPY 581/591 (Image-iT Lipid Peroxidation Kit, C10445, Invitrogen) for 30 min at 37 °C, followed by a single wash with PBS. Lipid ROS generation was assessed by fluorescence microscopy using an ECHO Revolve microscope. The reduced (590 - red) to oxidized (510-green) ratio was calculated to determine the lipid peroxidation.

### Labile iron pool assay and flow cytometry

The labile iron pool in the cells was analyzed by staining the cells with BioTracker Far-red Labile Fe^2+^ Dye (Sigma-Aldrich, SCT037). Briefly, the cells were treated with 5 μM of Far-red dye in serum-free media and incubated for 1 hour at 37 °C. After staining, the cells were washed once with PBS and detached from the plate using 0.25% trypsin EDTA. The cells were further centrifuged at 300g for 5 mins and resuspended in fresh PBS. Finally, the cells were filtered through a cell strainer (70 µm nylon mesh) and analyzed using a flow cytometer. The fluorescence signal was recorded using the APC filter.

### Total iron determination

The total iron content of cell lysates was assessed using the iron assay kit (ab83366, Abcam). Samples were prepared according to the manufacturer’s instructions and measured on a plate reader.

### Cell viability assay

Cell viability after RSL3 treatment was analyzed using the CellTiter-Glo® luminescent cell viability assay (Promega), which records luminescence on a plate reader.

### Western analysis

Protein expressions were detected by immunoblot as previously described. Briefly, cells were lysed using RIPA buffer (Sigma) and protease inhibitor cocktail (PIC, Sigma). Subsequently, total protein was quantified by BCA Protein Assay (Pierce), and equal amounts of protein were loaded onto a 4 to 20% Criterion TGX Precast Protein Gel (Bio-Rad). Proteins were transferred onto PVDF membrane and probed for FTH1 (Cell Signaling Technology, 1:1000, 4393S), beta-actin (Sigma, 1:1000, A5441), FTL (Abcam, 1:1000, ab69090), CD63 (Invitrogen, 1:250, 1062D), IRP2 (cell Signaling Technology;1:1000, #37135), FPN (alpha diagnostic, 1:1000, MTP11-S) . A corresponding secondary antibody conjugated to HRP was used (1:5000, GE Amersham), and the bands were visualized using ECL reagents (PerkinElmer) on an Amersham Imager 600 (GE Amersham).

### Fe57 treatment and ICP-MS

The cells were treated with 100uM ^57^FeCl_3_ (supplemented with 10uM sodium ascorbate) in serum-free media overnight. Now loaded with isotopically labeled iron, the cells were washed with PBS and cultured in fresh, serum-free media. After 24 hours, the cell lysates and the media were collected. The lysates and media were prepared by nitric acid digestion for ICP-MS (Inductively Coupled Plasma Mass Spectrometry) analysis, and the amount of ^57^Fe was quantified. Concentrations of ^57^Fe were normalized to lysate protein concentration to account for slight variations in seeded cell number.

### Exosome isolation

The exosome isolation from the cell culture media was completed as described previously [23, 24]. Briefly, the conditioned media were collected and centrifuged at (i) 300g for 10 min to remove the cell pellets, and (ii) the supernatant was centrifuged at 2000g for 10 min to remove the dead cell pellet. The resultant supernatant was centrifuged at 4000g for 30 min to remove the cell debris pellet. The resultant supernatant was collected in a 100K filter (Amicron Ultra -15, Centrifugal filters, Merck Millipore Ltd) to concentrate the media. The concentrated media were collected and centrifuged at 100,000g for 70 min (Beckman coulter, TLA 100.3). The supernatant was discarded, and the pellet was washed with PBS and again centrifuged at 100,000g for 70 min with the resultant exosome pellet used for downstream processing.

### Exosome inhibition studies

GW4869 (N, N’-Bis[4-(4,5-dihydro-1H-imidazol-2-yl) phenyl]-3,3’-p-phenylene-bis-acrylamide dihydrochloride) is the most widely used pharmacological agent to block EVs generation and reduce EVs release by neutral sphingomyelinase (nSMase). The nSMase activity is important for creating the large lipid raft domains involved in EVs shedding, as a result, inhibition of nSMase reduces the release of EVs from the cells. GW4869 was dissolved in DMSO and treated to the cells at a concentration of 30 μM for 24 hours to induce inhibition of exosome release. This dose of GW487 was chosen because this is the highest dose that can be used for 24 hours without inducing cytotoxicity [25] . Hepcidin (MedChemExpress HY-P4373) was added to the GBM cells to inhibit FPN expression at a concentration of 4 μM.

### TCGA Analysis

The correlation graph between CD68 and GPX4 was generated using the GLIOVIS (https://gliovis.bioinfo.cnio.es) data visualization tools. The adult TCGA GBM_LGG database was used for analysis and Pearson correlation was used for statistics.

### Statistics

All data were expressed as mean ± SD and the statistical analysis was done using GraphPad Prism 9. Students’ unpaired t-tests were used to compare the two groups. One-way ANOVA followed by Tukey’s post hoc analysis test was used to detect statistical significance (p < 0.05) between the multiple groups.

## Results

### Macrophage iron metabolism is sexually dimorphic

To determine whether sex influences macrophage iron handling, we isolated bone marrow-derived monocytes from C57BL/6 mice. After the differentiation of monocytes into macrophages in the presence of m-CSF, we examined the expression of L-ferritin (FTL), a protein essential for cellular iron storage. Our results demonstrated that the baseline expression of FTL was significantly higher in male BMDM than in females (**Fig. 1A, B**). The expression of FTL is directly proportional to the amount of stored iron; therefore, we treated the macrophages with two iron sources, ferric ammonium citrate (FAC) and ferumoxytol, which resulted in an increased expression of FTL. To understand whether the higher basal expression of FTL in male BMDMs corresponds to higher metabolically active iron, we measured the labile iron pool, where, similar to ferritin, male macrophages had significantly higher labile iron pool (**Fig. 1C**). Next, we hypothesized that male macrophages having higher stored iron and labile iron pool is a result of higher iron uptake. To address this, we treated the macrophages with serum-free media containing 100 μM ^57^Fe, a stable isotope of iron. After 24 hours, we measured the ^57^Fe in the macrophage lysates by ICP-MS. The analysis showed that male BMDMs took up significantly more ^57^Fe than female BMDMs (**Fig. 1D**). However, we did not see any sex difference in iron release from BMDMs (**Fig. 1E**). To mimic tumor-associated macrophages, we checked the iron release in BMDMs grown in GL261 conditioned media. The TAMs had a higher iron release than naïve macrophages, but there were no sex biased differences in iron release (**Fig 1E**). Next, to understand how macrophage iron metabolism is altered in the presence of GBM cells, we grew the BMDMs with GL261 cells in a transwell coculture system. Immunoblot analysis of macrophages after coculture with GL261 cells for 24 hours showed that the macrophages showed elevated FTH1 and FTL expression when in coculture with GL261 cells but lost their sex biased differences (**Fig. 1F, G, H**). Interestingly, there is a switch in sex biases for transferrin receptor expression when the macrophages are cocultured with GBM cells (**Fig. 1F, I**).

**Figure 1:**
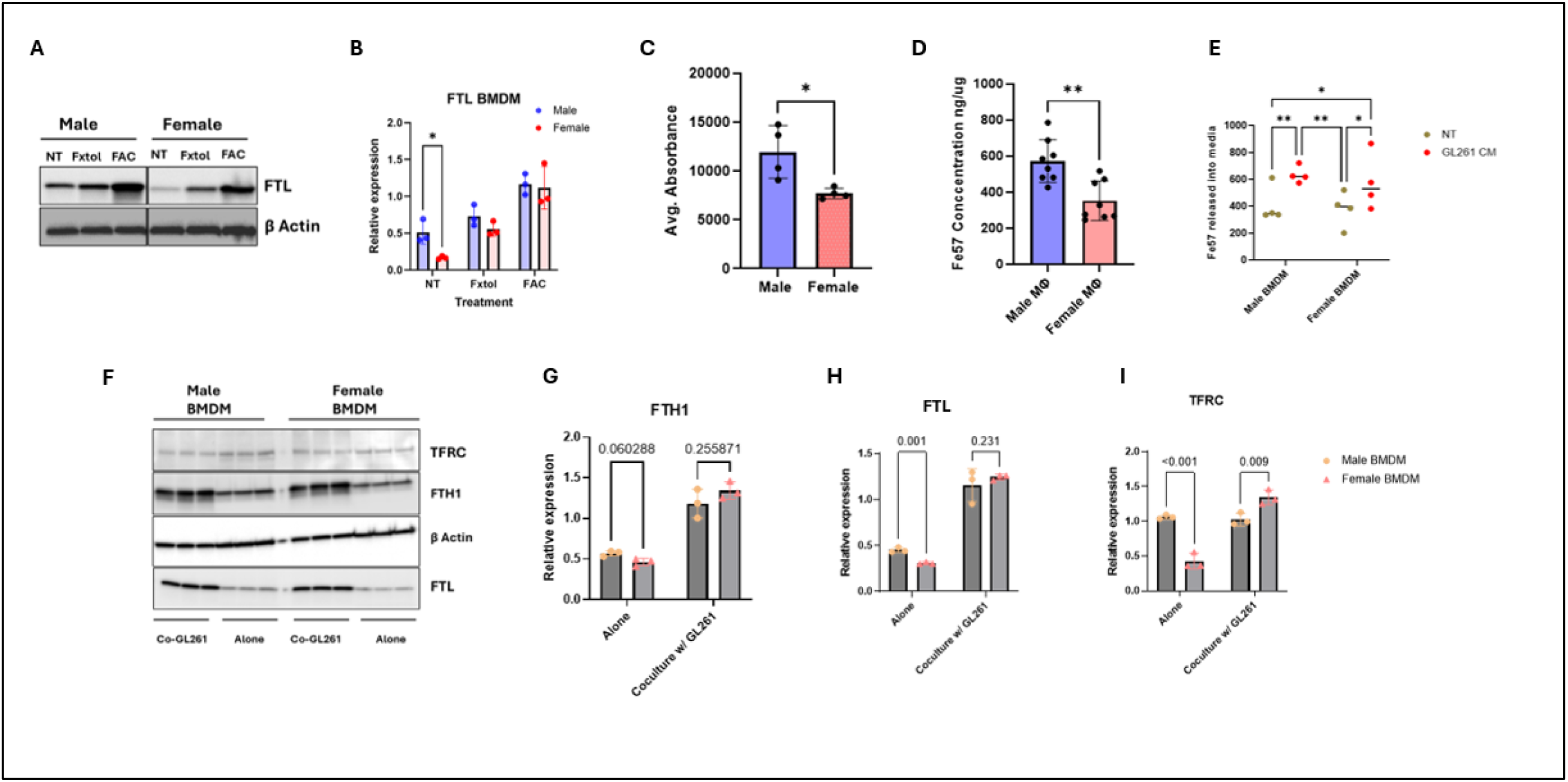
**(A)** Immunoblot for FTL expression in BMDM cells showing male BMDMs have a higher expression of FTL than females. Ferumoxutol (10 μM) and ferric ammonium citrate (100 μM) were treated to indicate an increase in FTL levels after iron loading. Beta-Actin was used as loading control. **(B)** graphical representation of immunoblot data. (n=3, unpaired t-test for NT male and female) **(C)** Labile iron pool measurement in BMDM showing male macrophages have higher labile iron (Fe2+) than females: cells were stained with BioTracker Far-red Labile Fe2+ Dye and analyzed in a flow cytometer. (n = 4, unpaired t test) **(D)** Iron uptake by BMDM was determined by measuring ^57^Fe uptake by ICP-MS. Male BMDMs showed higher 57Fe uptake than females. (n = 8, unpaired t test) **(E)** Fe57 release measurement from BMDM after treatment with GL261 conditioned media. GL261 conditioned media induced increased ^57^Fe release in both male and female BMDMs. (n = 4, two-way ANOVA). **(F)** Immunoblot for FTL, FTH1 and TFRC expression in macrophages cultured alone or with GL261 cells. **(G, H, I)** graphical representation of immunoblot data (n=3, one way ANOVA)

### TAMs induce a reduction of cellular iron and ferritin in GBM cells

Next, we hypothesized that TAMs could affect the iron metabolism of GL261 cells in a sex biased manner. To test this, we co-cultured GL261 with male and female BMDMs in a transwell coculture system. After 24 hours of coculture, we assessed the iron status of GBM cells by quantifying key cellular iron indicators, including ferritin, ferroportin (FPN), the Labile iron pool (Fe2+), and total cellular iron (Fe3+ and Fe2+). We observed a significant decrease in cellular FTH1 in GL261 cells when cocultured with BMDMs, with a notable sex bias, where the reduction in FTH1 in cells cocultured with male BMDMs was the most pronounced (**Fig. 2A, B**). Next, we checked the expression of FPN via immunoblotting and observed no alternations in FPN expression in GL261 cells after coculture male and female BMDM (**Fig. 2C**). Similarly, we did not see any difference in the cellular labile iron pool in cocultured GBM cells and GBM cells grown separately (**Fig. 2D**). Then, we measured the total iron, which includes both the labile iron pool (Fe2+) and protein-bound (Fe3+) iron, where we observed that coculture with BMDM significantly reduced the total iron in GL261 cells (**Fig 2E**). There was a sex-biased trend where GL261 cells cocultured with male macrophages had a higher reduction in cellular iron.

**Fig 2.**
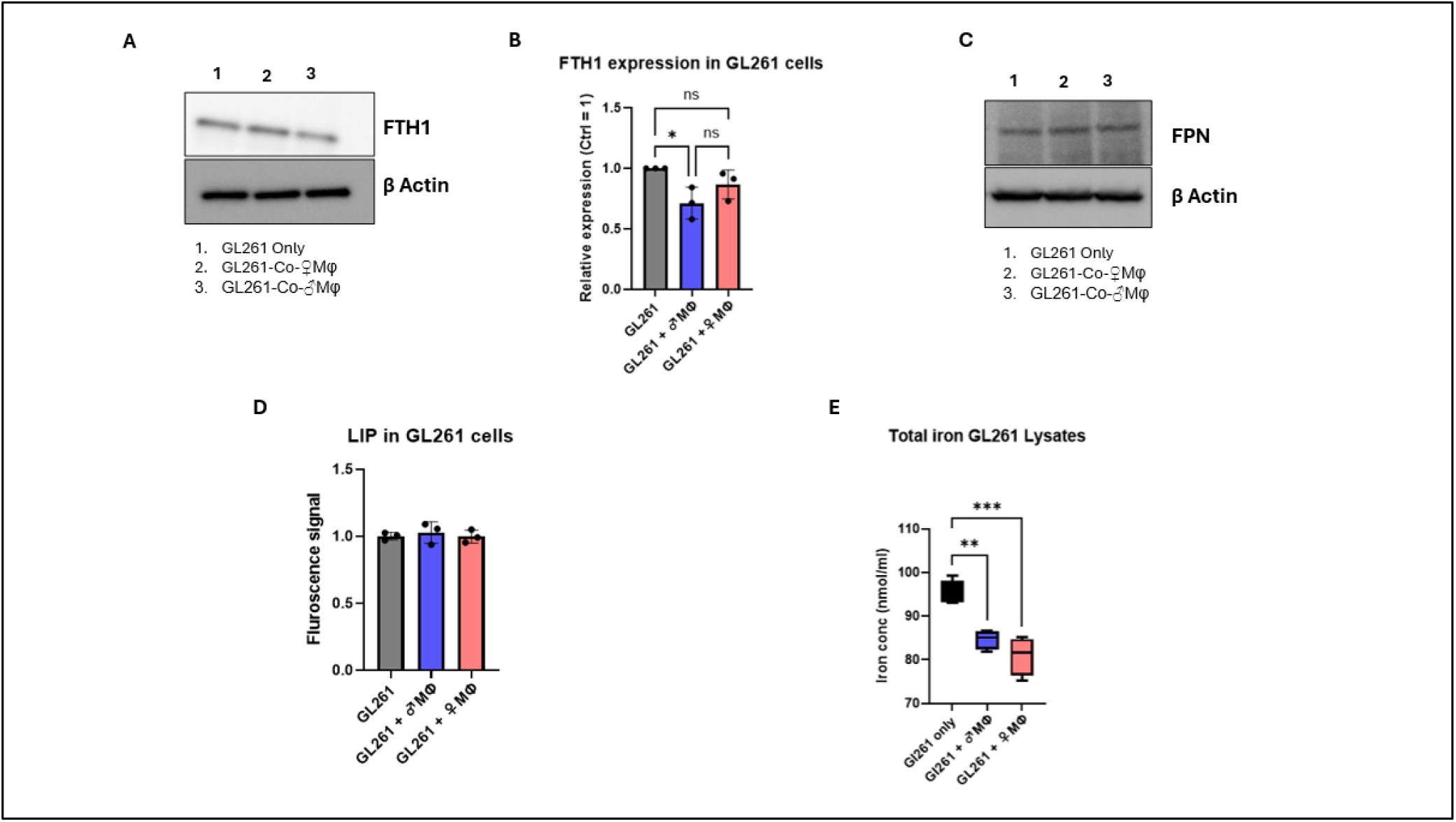
**(A)** Immunoblot showing a reduction in cellular FTH1 levels in GL261 cells that were cocultured with male BMDMs. Beta-Actin is used as loading control. **B:** graphical representation of immunoblot data. (n=3, One-way ANOVA) **(B)** Immunoblot showing no changes in the expression of FPN in GL261 cells cultured alone or in coculture with BMDM. Beta-Actin is used as loading control. **(C)** Ferro-far red assay showing no significant changes in cellular labile iron pool after coculture with BMDM. (n = 3, one-way ANOVA) **(D)** Cellular iron measurement assay showing a decrease in total cellular iron (Fe2+ and Fe3+) in GL261 cells after coculture with BMDM. (n=3, One-way ANOVA)

### TAMs induce the release of ferritin-bound iron from GBM cells

To understand how the GBM cells in coculture with BMDMs had reduced cellular ferritin and total iron, we hypothesized that TAMs induce the release of iron-bound ferritin from GL261 cells. To address this hypothesis, we cocultured the GL261 cells with male or female BMDMs for 24 hours to induce TAM phenotype in macrophages. Then, we removed the upper insert containing macrophages and added fresh serum-free media to the GL261 cells. Next, we collected the media from the TAM primed GL261 cells after 24 hours, which contained secreted proteins (**Fig. 3A**). Immunoblotting analysis showed that GL261 cells in coculture with BMDMs had a significantly higher FTH1 secretion in the media, with a clear male sex bias (**Fig 3B, C**).

**Fig 3:**
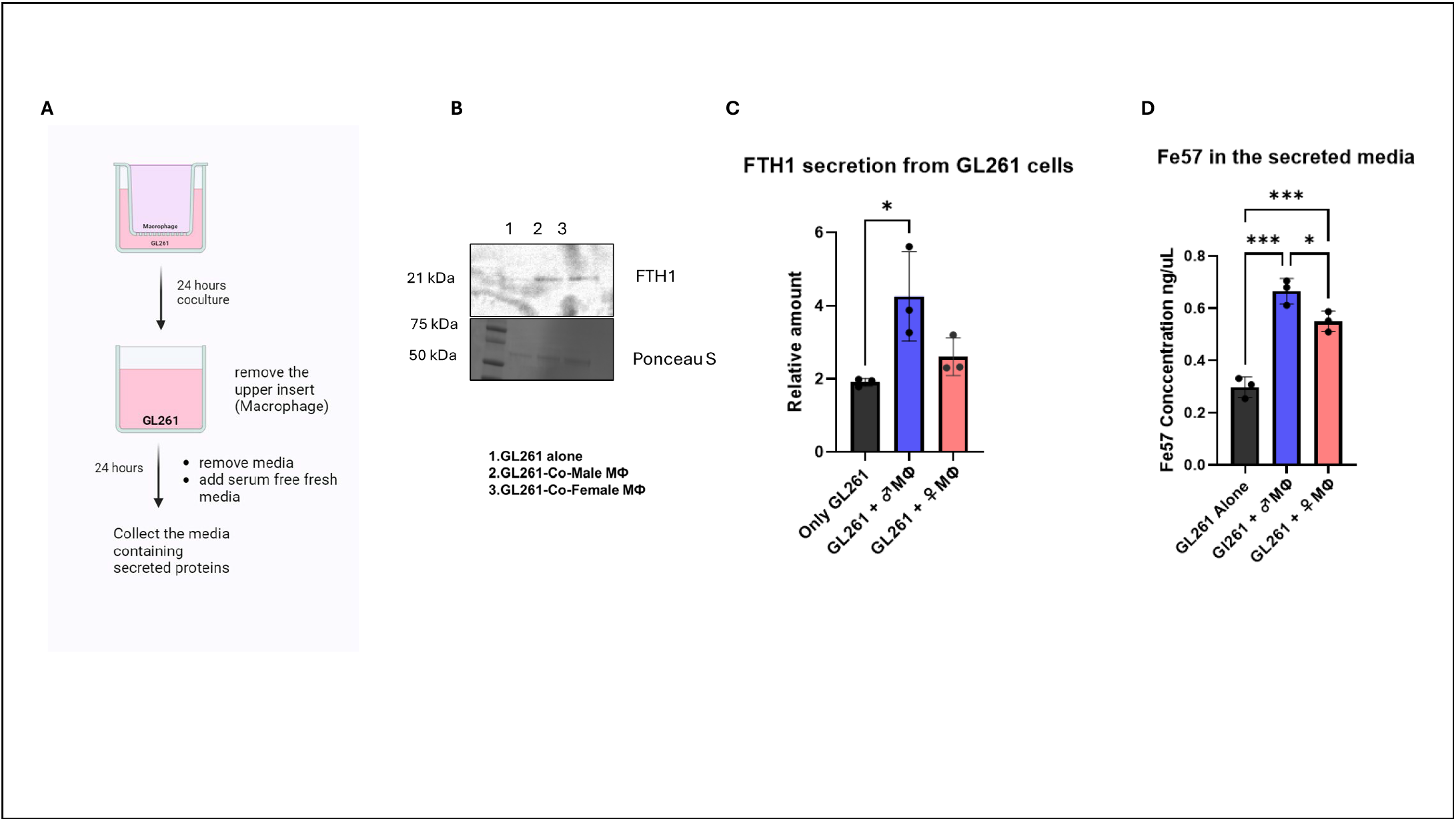
**(A)** Schematic diagram for measuring secreted FTH1 from GL261 cells. **(B)** Immunoblot showing higher FTH1 in the secreted media of the GL261 cells that were in coculture with male BMDM. Ponceau staining was done to use total protein as a loading control. **(C)** graphical representation of immunoblot data. (n=3, One-way ANOVA) **(D)** Fe57 release from loaded GL261 cells into the coculture media shows that BMDMs induce iron release from GBM cells in a sex biassed manner. (n=3, One-way ANOVA)

Next, to determine whether FTH1 secretion correlated with iron release, we used ^57^Fe-loaded GL261 cells and repeated the experiment. We observed that the ^57^Fe secretion was significantly increased in TAM-cocultured GL261 cells compared to GL261 cells cultured separately, with a clear male bias where male TAMs induced a higher iron release in GL261 cells than female TAMs (**Fig. 3D**).

### IRP2-regulated CD63-positive exosomes are involved in the release of ferritin

To understand the mechanism by which TAMs induce FTH1 secretion from GBM cells, we used immunoblotting to examine the cellular expression of Iron Regulatory Protein 2 (IRP2) in GBM cells cultured alone or in coculture with TAMs. We observed that GBM cells cocultured with male TAMs had a significant reduction in cellular IRP2 (**Figure 4A, B**). The IRP2 expression in GBM cells cocultured with female tumor-associated macrophages (TAMs) showed a lower trend compared to GBM cells alone. IRP2 impacts cellular iron uptake and release by post-transcriptionally regulating the expression of several key proteins involved in iron homeostasis. One such protein is CD63, which is involved in the biogenesis of exosomes. Thus, we checked the cellular expression of CD63 in the GBM cells. We observed that GL261 cells co-cultured with male TAMs had an increased expression of cellular CD63. (**Figure 4C, D**). We isolated the exosomes from the GL261-secreted media by sequential ultracentrifugation. We confirmed the isolation of exosomes by checking the expression of CD63. Finally, we observed that the exosomes secreted from GL261 cells cocultured with BMDMs had increased FTH1 and CD63 protein levels (**Figure 4E, F**).

**Fig 4.**
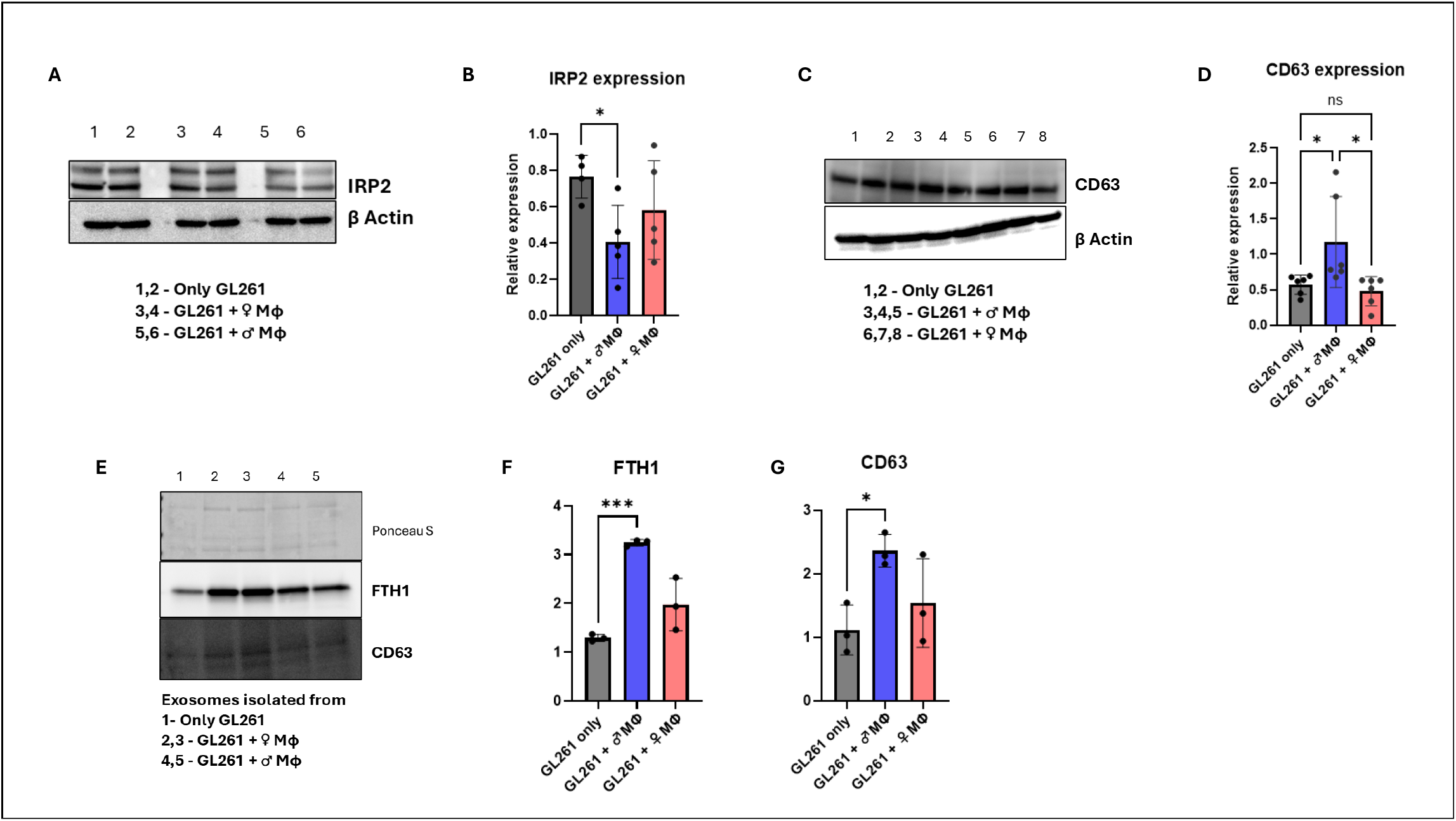
**(A)** Immunoblot showing a reduction in cellular IRP2 levels in GL261 cells that were cocultured with male BMDMs. Beta-Actin is used as loading control. **B:** graphical representation of immunoblot data. (n=4, One-way ANOVA) **(C)** Immunoblot showing an increase in cellular CD63 levels in GL261 cells that were cocultured with male BMDMs. Beta-Actin is used as loading control. **D:** graphical representation of immunoblot data. (n=6, One-way ANOVA) **(E)** FTH1 and CD63 protein expression in the **exosomes** isolated from the GL261 secreted media. The exosomes secreted from GL261 cells that were in coculture with male BMDMs showed increased CD63 and FTH1 levels. **(F, G)** graphical representation of immunoblot data. (n=3, One-way ANOVA)

### TAMs protect GBM cells from RSL3-induced ferroptosis

A decrease in cellular iron can make cancer cells more resistant to iron-dependent cell death. To analyze whether TAM infiltration had any impact on ferroptosis-related genes in glioblastoma, we used bioinformatics analysis from TCGA to correlate the expression of macrophage-specific marker CD68 with GPX4, a protein directly involved in ferroptosis suppression in cells. We observed a positive correlation between CD68, representing tumor-infiltrating macrophages, and GPX4, indicating that GBM tissues with higher TAMs could be more resistant to ferroptotic stress (**Fig. 5A**). To validate this hypothesis in vitro, we used a transwell coculture system with GL261 cells with TAMs (male and female). Next, to analyze the impact of TAMs on tumor cell ferroptosis, we used a transwell co-culture system in which GL261 cells were grown on the bottom and the macrophages on the top. Subsequently, 3 μM of RSL3 was added to the wells for 24 hrs. RSL3 induces ferroptotic stress by inactivating glutathione peroxidase 4 (GPX4). After 24 hours of treatment, the upper insert was detached, and the viability of the GBM cells in the lower chamber was determined using Cell Titer Glo. The viability assay showed that the GBM cells in coculture with macrophages had significantly more viable cells after RSL3 treatment (**Fig. 5B**). To confirm that the cell death was specifically due to ferroptosis, we added liproxstatin-1(Lip), a potent inhibitor of ferroptosis, and observed that adding Lip restored the cell viability even after the treatment with RSL3. Lipid peroxidation is a classic marker for ferroptosis. Hence, we investigated the level of lipid peroxidation in GBM cells using Bodipy staining. Using a lower dose of RSL3 (1uM) to induce non-lethal ferroptotic stress, we found a significant decrease in the 590/510 (non-peroxidized lipids/peroxidized lipids) ratio, indicating higher lipid peroxidation in non-cultured GBM cells treated with RSL3. However, the GBM cells in co-culture showed a significant rescue in the 590/510 ratio, indicating that macrophages reduce the amount of lipid peroxidation in GBM cells after RSL3 treatment (**Fig. 5C**). We used both male and female macrophages to determine whether there were any potential sex differences. However, despite some sex-biased trends in lipid peroxidation amounts, they failed to reach significance. Finally, we observed that RSL3 treatment increased the LIP in GL261 cells by FerroFarRed assay (**Fig. 5D**). The RSL3-treated GL261 cells that were in coculture with macrophages had significantly reduced LIP than the ones cultured alone.

**Fig 5:**
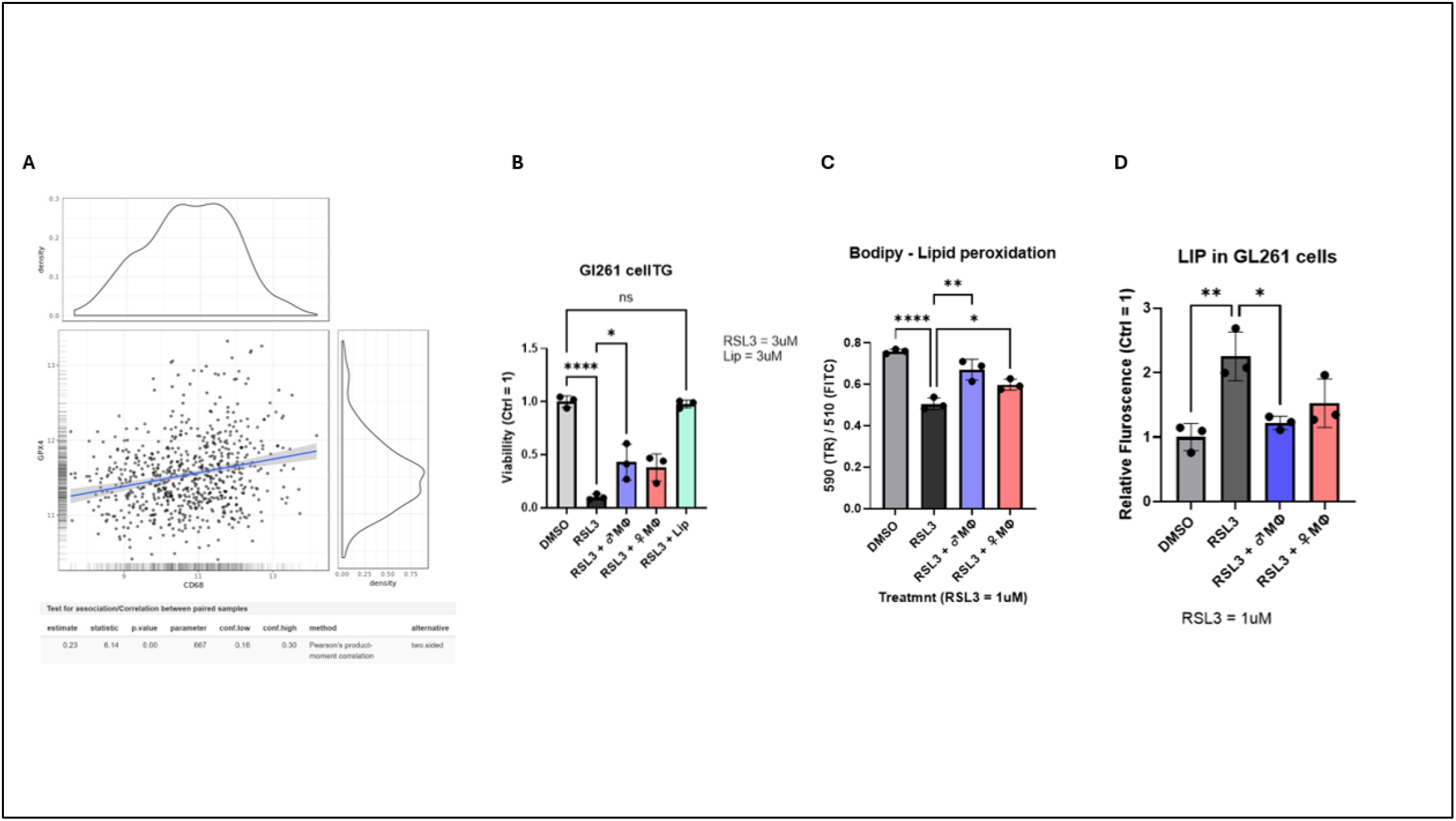
**(A)** Bioinformatics analysis showing a positive correlation between CD68 and GPX4 in GBM tissue from patients. (TCGA, Gliovis database) CD68 is used as a TAM marker, and GPX4 as a marker for the suppression of ferroptosis. A positive correlation indicates that GBM tissues with higher macrophage infiltration can have higher resistance to ferroptotic stress. **(B)** Cell Titer Glo viability assay showing that GL261 cells in coculture with BMDMs show resistance to cell death after RSL3-induced ferroptotic stress. (RSL3 = 3uM, Lip = 3uM) (n=3, One-way ANOVA). Liproxastatin 1(Lip) is used as an inhibitor of ferroptosis. **(C)** Bodipy staining to determine lipid peroxidation after RSL3 treatment indicates that GL261 cells in coculture with BMDM have lower lipid peroxidation. Lipid peroxidation shifts the fluorescence emission peaks from 590 nm to 510 nm after Bodipy staining. (RSL3 = 1uM) (n=3, One-way ANOVA) **(D)** Labile iron pool measurement in GL261 cells showing coculture with BMDM reduces Fe2+ in GL261 cells under ferroptotic stress induced by RSL3. (RSL3 = 1uM) (n=3, One-way ANOVA)

### Ferroptosis resistance by exosome-mediated FTH1 release is independent of the Hepcidin/ ferroportin axis

To understand whether the exosome-mediated FTH1 release pathway is potentially involved in ferroptosis resistance during inflammation, we treated the co-culture with Hepcidin, a protein that is elevated during inflammation. Hepcidin is an inhibitor of Ferroportin (**Fig. 6A**) and it blocks FPN-mediated iron release. Hepcidin treatment (4uM) did not alter the ferroptotic resistance of GL261 cells. To confirm that the suppression of ferroptosis is regulated by exosome release during inflammation, we treated both GW4869 and Hepcidin to maximally inhibit iron export from the cell. GW4869 is a sphingomyelinase inhibitor that reduces exosome biogenesis (**Fig. 6B**). This treatment resulted in the loss of ferroptotic resistance in BMDM-cocultured GL261 cells after RSL3 treatment (**Fig. 6D**). Finally, we loaded the GL261 cells with ^57^Fe (100uM) and measured iron release after Hepcidin and Hepcidin + GW4869 treatment. Hepcidin treatment significantly decreased iron release compared to control. The greatest decrease in iron release was obtained when Hepcidin and GW4869 treatments were combined (**Fig. 6E**)

**Fig 6.**
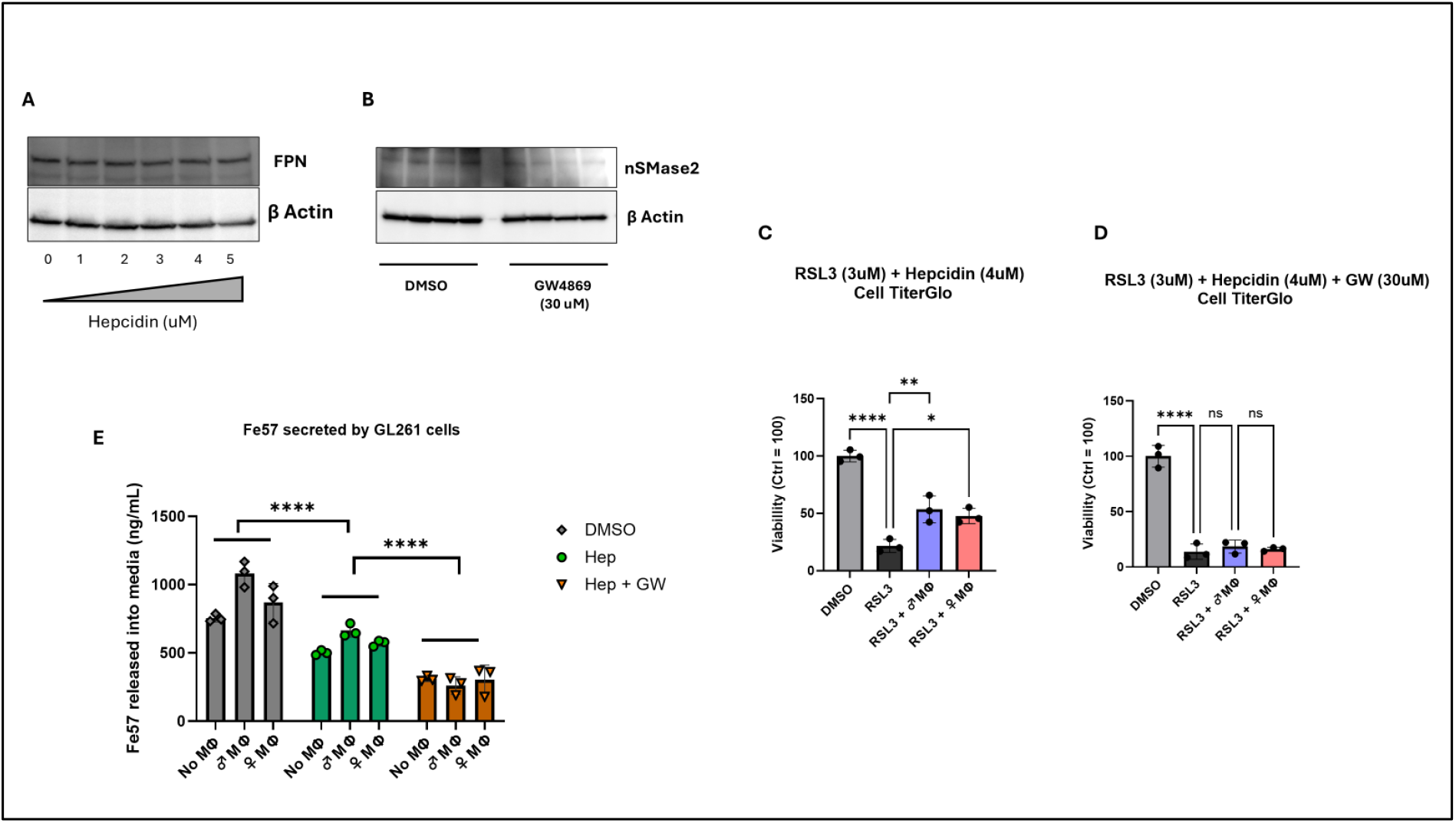
**(A)** Immunoblot showing dose-dependent inhibition of FPN by Hepcidin in GL261 cells and **(B)** inhibition of nSMase2, a key enzyme involved in exosome secretion by GW4869 (30uM) in GL261 cells. Beta-Actin is used as loading control. **(C)** Cell Titer Glo viability assay showing that GL261 cells in coculture with BMDMs show resistance to cell death even when FPN-mediated iron release is inhibited by Hepcidin after RSL3-induced ferroptotic stress. (RSL3 = 3uM, Hepcidin = 4uM)) (n=3, One-way ANOVA). **(D)** Cell Titer Glo viability assay showing that GL261 cells in coculture with BMDMs lose the resistance to cell death when both FPN and exosome-mediated iron release are inhibited by Hepcidin and GW4869 treatment after RSL3-induced ferroptotic stress. (RSL3 = 3uM, Hepcidin = 4uM, GW4869 = 30uM) (n=3, One-way ANOVA). **(E)** Fe57 release measurement from GL261 cells after treatment with DMSO (control), Hepcidin, and Hepcidin + GW4869. There is a decrease in ^57^Fe release with respect to the control after Hepcidin treatment, and a further decrease in ^57^Fe after Hepcidin + GW4869 treatment. (n=3)

## Discussion

Our data indicate that TAMs play a crucial role in regulating GBM iron metabolism and making it a sexually dimorphic disease. The TAMs regulate iron availability in the TME by acting as iron reservoirs that store excess iron in the tissue and distribute it when required [26]. By regulating the iron availability in the TME, TAMs affect tumor growth, immune infiltration, ferroptotic stress, and hypoxia-mediated angiogenesis [5, 27, 28]. Typically, high TAM infiltration is linked to worse prognosis and resistance to therapy, as it can promote tumor growth, metastasis, angiogenesis, and hypoxic signaling [29, 30]. There have been multiple reports of TAMs suppressing ferroptosis in cancer cells, especially in prostate cancer, cervical cancer, and fibrosarcoma. In this study, we investigate how TAMs influence the ferroptotic susceptibility of GBM cells, with a particular focus on sex-biased differences in iron metabolism. Ferroptosis is a type of iron-mediated cell death, where iron plays a pivotal role in catalyzing lipid peroxidation, a classic hallmark of ferroptosis via the Fenton reaction. Hence, cellular iron fluctuations in tumor cells can alter their susceptibility to ferroptotic stress. Previous publications have explored possible mechanisms by which TAMs influence tumor ferroptosis, primarily by modulating signaling pathways associated with lipid peroxidation, such as the LXRα/SCD1 pathway in prostate cancer and the ALOX15 pathway in cervical cancer [31, 32]. The only known mechanism of ferroptosis regulation by TAMs that directly involves iron regulatory pathways has been demonstrated in fibrosarcoma, where TAMs induce the upregulation of ceruloplasmin in cancer cells, resulting in iron export via FPN and resistance to ferroptosis [14]. However, due to the high proinflammatory signaling in the tumor microenvironment (TME) of several cancers, such as GBM, there is an upregulation of hepcidin, which can inhibit FPN-mediated iron release [33]. We propose a novel alternate mechanism by which TAMs induce resistance to ferroptotic stress in GBM cells, especially during high inflammation in the TME. Previous work from our laboratory has demonstrated that H-ferritin (FTH1) functions as an iron-delivery protein and is involved in iron translocation[34]. FTH1-bound iron release from cells can serve as an alternative to ferroportin-mediated iron release, particularly during periods of inflammation. Further past research from our laboratory has shown that exosome-mediated iron release is a noncanonical pathway for iron release, which is ferroportin-independent but regulated by the IRP/CD63 pathway [24, 35]. In this study, we address how TAMs regulate the iron status and ferroptotic susceptibility of GBM cells.

It is already well-established that systemic iron metabolism is sexually dimorphic. Males have higher serum iron, hemoglobin, and hepcidin. However, at the tissue and cellular levels, few studies have established clear sex differences in iron homeostasis. To understand whether macrophage iron metabolism is sex-biased, we used male and female bone marrow-derived macrophages (BMDMs). We observed distinct sex differences in the macrophage iron metabolism. Male macrophages showed higher iron storage (FTL expression) and a higher labile iron pool. These differences in protein-bound and free iron levels between male and female BMDMs are a result of sex-biased iron uptake, as demonstrated by the ^57^Fe uptake experiments. These data suggest that macrophage iron metabolism is sex-biased, with male macrophages exhibiting higher iron uptake, storage, and labile iron. The trends observed on a systemic level, i.e., males have a higher iron content in their serum and tissue than females, also hold true at a cellular level in terms of macrophage-specific iron metabolism. However, the innate sex biases seem to alter when the macrophages are grown in the presence of GL261 cells. The higher expression in male macrophages of FTH1 and FTL is lost when in coculture with GBM cells. Similarly, the TFRC expression, which is higher in male macrophages when grown alone, changes to higher in females after coculture with GL261 cells. The data indicate that there exist sex specific biases in macrophage iron metabolism in the TME, which is different from the sex biases that are observed in normal macrophages.

In glioblastoma, TAMs can make up as much as 50 percent by volume of the total tumor and are the most abundant cell type present in the TME. However, the impact of TAM infiltration on ferroptosis-related genes is still unknown. We performed bioinformatic analysis on the GBM TCGA database to correlate GPX4 expression with CD68, a marker for macrophages, across the entire tumor transcriptome. GPX4 is a selenoprotein and antioxidant enzyme that plays a crucial role in protecting cells from lipid peroxidation and ferroptosis. We observed a positive correlation between CD68 and GPX4 expression in clinical data obtained from TCGA, suggesting that higher TAM infiltration may contribute to resistance to ferroptosis in GBM tissue. In an in vitro coculture system, we observed a similar effect, where the GL261 cells cocultured with BMDMs showed resistance to RSL3-induced ferroptotic stress both by viability studies and lipid peroxidation. Lipid peroxidation is a hallmark of ferroptosis, where cells undergoing ferroptosis show a substantially higher amount of lipid peroxidation. These results are consistent with previous publications showing that tumor-associated macrophages can confer ferroptotic stress resistance to tumor cells.

To understand the mechanism by which TAMs induce ferroptosis resistance in GBM cells, we used our coculture system to determine how TAMs affected the overall iron status of the GBM cells. GL261 cells in coculture with TAMs exhibited a reduction in overall cellular FTH1 with a clear sex bias, where male TAMs induced a higher decrease in FTH1 than females. FTH1 can act as both an iron storage and an iron delivery protein in the cell, binding to Fe^3+^ iron. However, we observed that TAMs don’t alter the labile iron pool (Fe^2+^) in the GBM cells under normal conditions, indicating that only the protein-bound Fe^3+^ iron is affected. Similarly, we didn’t observe any changes in the FPN expression, indicating that BMDMs do induce changes in the FPN mediated Fe^2+^ iron release. The total cellular iron measurement in the GBM cells confirmed that GL261 cells cocultured with TAMs exhibited a significant reduction in total cellular iron (Fe^2+^ + Fe^3+^). There was also a sex bias in total cellular iron, where GL261 cells cocultured with male TAMs exhibited a higher iron reduction. To understand how cellular FTH1 levels were reduced, we measured the secreted FTH1 levels from the TAM-primed GL261 cells and found that GL261 cells cocultured with TAMs secreted more FTH1 than those cultured alone. These data align with the decrease in cellular FTH1 levels. Next, we asked whether this increase in FTH1 secretion correlated with an increase in iron release from the GL261 cells. ^57^Fe experiments demonstrated that GBM cells in coculture with TAMs secreted more iron than non-cocultured cells, with a clear male bias. These data indicate that TAMs induce FTH1-bound iron release from GBM cells, representing a major pathway for iron release. This reduction in cellular iron contributes to resistance to ferroptosis in GBM cells.

To understand how TAMs induce FTH1 release, we examined the potential pathway for FTH1 secretion. Coculture of GL261 cells with TAMs caused a decrease in overall cellular IRP2 levels in GBM cells. There was a clear sex bias, with only the GBM cells cultured with male TAMs showing significance; females exhibited only a trend. IRP2 can directly regulate exosome biogenesis via CD63, which has an IRE sequence in the 5’-untranslated region (UTR) of its mRNA, and only GBM cells cocultured with male tumor-associated macrophages (TAMs) showed an increase in cellular CD63. Because CD63 is a major protein involved in exosome biogenesis, we hypothesized that the cells secreted iron-bound FTH1 via CD63-positive exosomes. Hence, we isolated the exosomes from the GL261 secreted media by ultracentrifugation. The exosomes secreted from GL261 cells that were previously in coculture with TAMs showed higher FTH1 and CD63 expression with a strong male bias.

Previous work from our laboratory and others has shown that CD63-positive exosomes are involved in the cellular release of iron. This noncanonical way of iron release is independent of Hepcidin regulation [24]. Our data indicate that iron release occurs from the GL261 cells via both ferroportin and exosomes. Thus, release via exosomes can be a potential pathway by which GBM cells escape ferroptosis stress during inflammation. Inflammation results in increased secretion of hepcidin by astrocytes into the TME, which will block iron release by FPN, resulting in higher iron available for the Fenton reaction and higher ferroptosis [36]. Moreover, ferroptotic resistance could be conferred by the TAMs in the presence of hepcidin by promoting GL261 iron release via exosomes containing iron-bound FTH1. Finally, to confirm that exosome release is the major pathway that confers ferroptosis resistance, we treated the coculture with GW4869, an inhibitor of exosome biogenesis, which resulted in cell death, indicating that the ability of TAMs to protect the GL261 cells was eliminated when exosome release was blocked in the GL216 cells. ^57^Fe experiments confirmed that Hepcidin treatment resulted in a significant decrease in iron release, and the co-treatment of GW and Hepcidin caused an even higher reduction in iron release.

In summary, this study reveals that FTH1 release via CD63-positive exosomes is a significant mechanism via which TAMs confer ferroptotic resistance to GBM cells. There appears to be a moderate sex bias in this process that favors males. This cell culture-based study may provide insights into the clinical observations that male GBM is more resistant to standard therapy because of the decrease in ferroptosis within the tumor. We did observe that female TAMs can confer some resistance to GL261 cells, but the process seems to have less reliance on CD63-positive exosome-mediated FTH1 release.

## Acknowledgments

We thank Dr Maggie Wang, Ph.D., Laboratory for Isotopes and Metals in the EESI, Penn State University, State College, for the ICP-MS. Fig. 3A was created with BioRender.com. JRC would like to acknowledge funding received from the NCI-NIH (5P01CA245705-05).

## Authors contribution

APS and JRC conceived the study. APS and KP performed the experiments. BSW supplied reagents and mice. APS, KP, and GS analyzed the data. APS and JRC drafted the manuscript. All authors read, edited, and approved the final version of the manuscript.

## Data availability

No datasets were generated during the current study.

